# Ecological and socioeconomic factors associated with globally reported tick-borne viruses

**DOI:** 10.1101/2024.10.11.617937

**Authors:** Samantha Sambado, Sadie J Ryan

**Author notes:** **Corresponding author**: Samantha Sambado, MS, MA; University of California, Santa Barbara. Marine Science Institute, Bldg. 520. Santa Barbara, California 93106-6150. **Conflict of interest**: The authors declare they have no conflicts of interests.

## Abstract

**Background:** Public health resources are often allocated based on reported disease cases. However, for lesser-known infectious diseases, such as tick-borne viruses, disease risk reporting should account for more than just the biology of the disease and include mediating factors such as socioeconomics which can determine if an infection gets reported.

**Objectives:** We aim to identify country-level ecological and socioeconomic factors important to reporting tick-borne viruses and examine whether countries with more economic resources have a higher likelihood of reporting resource intensive incidences. Our study goals are to determine potential country-level interventions that could enhance recognition of and reduce the health burden associated with tick-borne viruses.

**Methods:** We apply machine learning to the most comprehensive tick-borne virus database, ZOVER, with a curated global trait matrix of 23 environmental and socioeconomic predictors.

**Results:** We identified socioeconomic factors driving reported tick-borne viruses captured in the database at a country level. Countries that were more likely to report tick-borne viruses had a lower Gini Index (i.e., countries with less inequalities such as Nordic countries), increased dollars spent on pesticide imports, and had institutions (i.e., IVSA chapter) or individuals with agricultural, forestry, or veterinary knowledge (i.e., % of tertiary grads) present. Additional characteristics included countries with a lower percent of population exposed to conflict also had a higher probability of reporting a tick-borne virus. As expected, broad environmental factors such as the Köeppen-Geiger climate classification zone was important and identified Mediterranean climate or humid subtropical climate as environmentally suitable zones for reported tick-borne viruses.

**Discussion:** For environmentally persistent pathogens, the role of ancillary factors mediating reporting must be considered for allocating resources to interventions. In addition, while direct interruption of transmission is important, socioeconomic interventions may be the greatest tool to reduce local disease burden.

## INTRODUCTION

Infectious disease occurs when people and pathogens align in space and time^1–3^. However, reporting an occurrence of an infectious disease involves more than just the biological aspects of the disease system^4–7^. From the initial infectious contact to the reporting of a disease case, several steps must be completed (Fig. 1). First, individuals need to be aware of and recognize the disease symptoms. If health care is accessible, they must have the resources to attain care.

**Figure 1.**
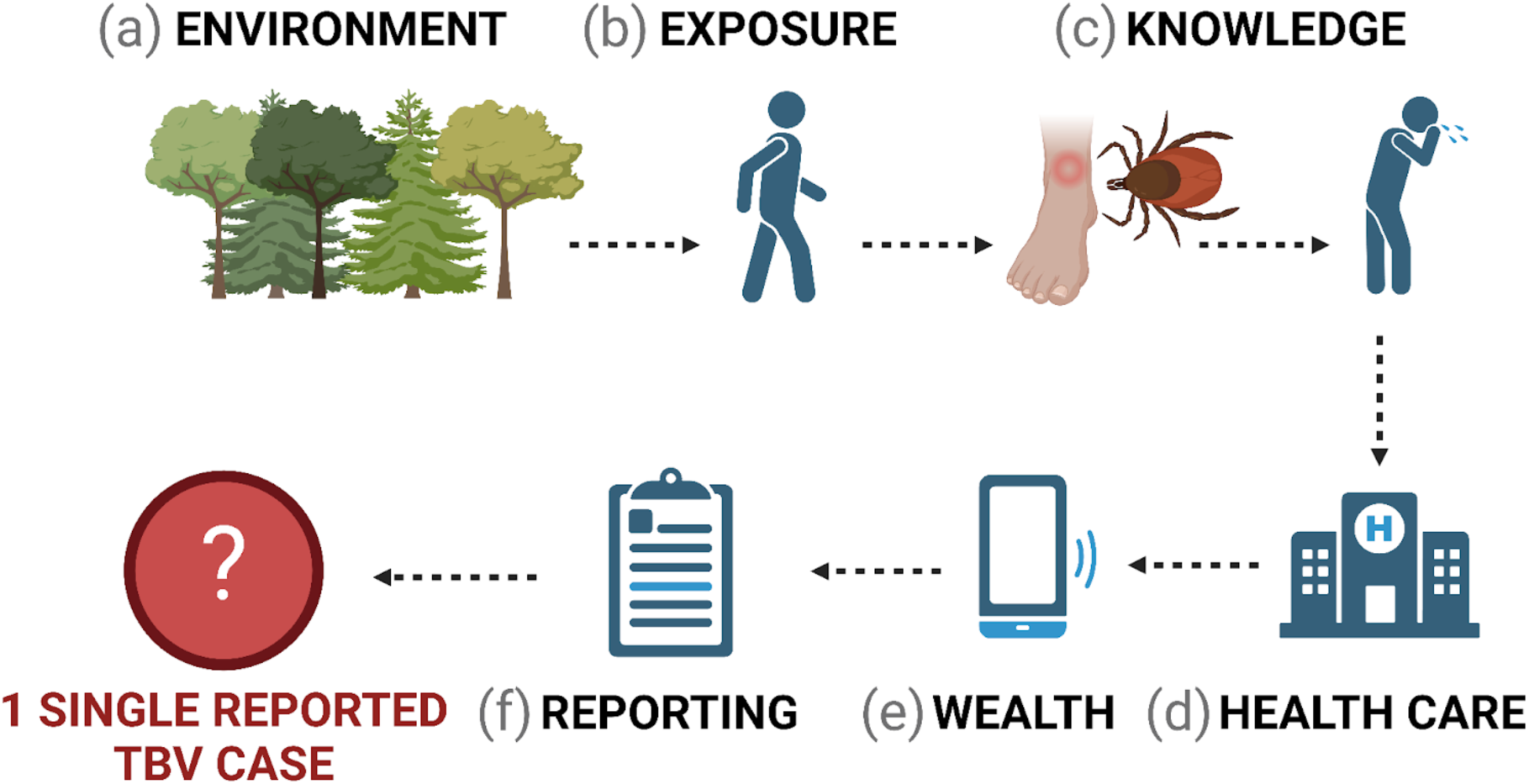
Illustration of the multi-tiered process involved in reporting a single tick-borne virus (TBV) case. It encompasses several critical factors: (**a**) a suitable environment for tick-borne pathogens; (**b**) human exposure to an infected tick; (**c**) awareness of ticks and associated disease symptoms (i.e., knowledge); (**d**) access to health care and (**e**) necessary resources for treatment (i.e., wealth); (**f**) a reporting system for documenting cases and estimating relative disease risk.

Following this, a systematic process must be in place to record and report the incidence, which is typically managed at the administrative level rather than by individuals. While the environmental suitability of a pathogen to persist in its natural cycles is undeniably important^8–10^, socioeconomic factors often play a significant role in determining which diseases are reported^4,5,11^. Reported case numbers, in turn, influence how public health resources are allocated.

The intensity of contact between people and pathogens are shaped by the characteristics of the surrounding landscape^6,10,12,13^. Risk of pathogen exposure can arise from various sources, including occupational hazards (e.g., individuals involved in agriculture or livestock management), residential environment (e.g., living near habitats conducive to pathogens like forests or animal aggregations), and recreational activities (e.g., walking through wooded areas)^4,14–17^. In contrast, while pathogen exposure might be lower in affluent urban environments, resources to manage potential exposures are typically more accessible compared to less urbanized communities^5,11,16,18^.

Infectious diseases impose a disproportionate health, and economic burden, on low-income countries^19–22^. The source of this disparity – whether due to biophysical features (such as increased biodiversity leading to higher pathogen diversity observed at lower latitudes) or reliance on natural resource-based activities (such as hunting, agriculture, or pastoralism for primary economic activity) – is not fully resolved for all diseases^5,12,19,20^. Nevertheless, key ecological insights offer a clearer understanding of certain diseases, especially those with well-documented life cycles and significant clinical impact, such as those with high morbidity and mortality^12,15,23^. For diseases with unclear transmission cycles or ambiguous symptoms, accurately assessing the true burden across all communities remains a significant challenge^15,18,24,25^.

Launched in 2022 by the World Health Organization, the Global Arbovirus Initiative aims to address the increasing risk of arbovirus epidemics, including tick-borne viruses (TBVs). Ticks transmit a larger range of pathogens (i.e., protozoa, rickettsiae, spirochetes and viruses) than any other arthropod group, representing a high burden of global economic impact due to their role in livestock, companion animal, and human diseases^15,24,26^. Tick-borne viruses, which are considered zoonotic infectious diseases, are reported to be increasing and spreading into new geographic regions at rates comparable to other zoonotic threats^15,24,27–29^. This reported increase may be influenced by advances in molecular techniques and potential study biases, rather than rising transmission rates^15,28,30–32^. Confirming a TBV often requires a multi-tiered approach, including accurate tick identification, recognition of noticeable symptoms, and reliable diagnostic tests^14,15,27,29,32^. While some tick-borne diseases, such as Lyme disease, Crimean-Congo hemorrhagic fever, and Tick-borne encephalitis are tracked by global surveillance platforms like WHO Disease Outbreak News, ProMed-mail, World Animal Health Information System, and National Immunization Technical Advisory Groups, this coverage does not reflect the full range of tick-borne diseases to which both urban and rural populations may be exposed^24^. Additionally, the knowledge bases for reporting and surveillance are still patchily distributed across geographies and stakeholders such as public health, veterinary, agricultural, and entomological agencies, as well as research hubs^26,33^. The high resource and knowledge requirements for reporting TBVs contributes to a greater likelihood of their detection in affluent communities, where such resources are more readily available, rather than in areas with the highest environmental and occupational risk^4,5,7^.

Addressing the impact of socioeconomic factors on treatment access and the prioritization of reported cases in public health decisions presents a crucial opportunity to develop strategies that can significantly alleviate the burden of disease^15,27,29,32^. Effective strategies often include socio-ecological levers, which focus on interrupting environmental exposure and reducing vulnerability^3–5^. These approaches have been successful in managing other infectious diseases with environmental transmission, such as mosquito control measures like bednets and water treatments^24,27,34^. However, for vectors that are relatively long-lived and less understood, similar strategies may not be effective for ticks and may need to be tailored to specific tick-borne diseases^29,34^. Nonetheless, it is essential to implement education campaigns, including increasing the popularity of community science programs, that empower patients and physicians with knowledge about TBVs^33^. Resources for effective tick control should be strategically designed at the residential level while implementing a standardized system for reporting and notifying about tick-borne viruses at local, regional, or national levels^33^.

Our study examines the environmental and socioeconomic factors driving reported TBVs at the country-level, aiming to identify potential interventions for enhancing recognition and reducing disease burden. We have compiled a database of 23 hypothesized drivers for each country and integrated it with the currently most comprehensive tick-virus database (ZOVER) to train our machine learning algorithm, boosted regression trees (BRTs). Generalized BRTs are well-suited for high-dimensional data and can effectively determine the relative importance of each driver on reported virus infected ticks^35–37^. This approach addresses some of the challenges in understanding TBV risks, which have been previously constrained by limited data availability and inadequate statistical tools^15,29^. This paper aims 1) to identify the key country-level environmental and socioeconomic characteristics associated with reported tick-borne viruses, and 2) to examine whether countries with greater economic resources have a higher likelihood of reporting these resource intensive incidences.

## Methods

### Data Curation

#### Reported Tick-borne viruses

To assess our key focus – reported tick-borne viruses at the country level – we use data from the most comprehensive current database on zoonotic and vector-borne viruses (ZOVER, http://www.mgs.ac.cn/ZOVER, accessed on 21 09 2024). Data on ZOVER were manually gathered from published literature and records from the GenBank database^30,38^. While ZOVER data was reported as infected ticks, we assume that if a country is screening ticks for a specific virus, it likely has the resources to report human cases of the same virus. We compiled a total of 4,213 unique records for the top tick-borne viruses of concern based on recent literature: African swine fever, Alkhumra hemorrhagic fever, Bhanja virus, Bourbon virus, Colorado tick fever virus, Crimean-Congo hemorrhagic fever, Deer tick virus, Heartland virus, Jingmen tick virus, Kyasanur forest disease virus, Louping ill virus, Lumpy skin disease, Nairobi sheep disease virus, Omsk hemorrhagic fever, Powassan virus, Sawgrass virus, Severe Fever with Thrombocytopenia Syndrome virus, Tick-borne encephalitis (Table 1)^14,24,28,31,39^.

**Table 1.** List of viruses included in this study including disease name, disease abbreviation, virus family, known vectors of virus, and number of PubMed citations per disease (accessed on 2024-08-09). Virus name and viral family come from NCBI Taxonomy Browser.

| Disease | Abbrev. | Viral family | Vector Genus | PubMed |
| --- | --- | --- | --- | --- |
| African swine fever virus | ASFV | Asfarviridae | *Ornithodoros* | 3,253 |
| Alkhumra hemorrhagic fever virus | AHFV | Flaviviridae | *Ornithodoros* | 33 |
| Bourbon virus | BRBV | Orthomyxo-<br>viridae | *Amblyomma*<br>*Haemaphysalis* | 43 |
| Colorado tick fever virus | CTFV | Spinareoviridae | *Dermacentor* | 190 |
| Crimean-Congo hemorrhagic fever virus | CCHFV | Nairoviridae | *Amblyomma*<br>*Hyalomma*<br>*Haemaphysalis*<br>*Rhipicephalus* | 1,882 |
| Deer tick virus | DTV | Flaviviridae | *Ixodes* | 1,590 |
| Heartland virus | HRTV | Phenuiviridae | *Amblyomma* | 111 |
| Jingmen tick virus | JMTV | Flaviviridae | *Rhipicephalus*<br>*Haemaphysalis*<br>*Ixodes* | 56 |
| Kyasanur forest disease virus | KFD | Flaviviridae | *Haemaphysalis* | 264 |
| Lumpy skin disease virus | LSDV | Poxviridae | *Hyalomma*<br>*Rhipicephalus* | 576 |
| Nairobi sheep disease virus | NSDV | Nairoviridae | *Haemaphysalis* | 131 |
| Powassan virus | POWV | Flaviviridae | *Ixodes*<br>*Dermacentor*<br>*Haemaphysalis* | 4,211 |
| Severe fever with thrombocytopenia syndrome virus | SFTF | Phenuiviridae | *Haemaphysalis* | 1,431 |
| Tick-borne encephalitis virus | TBEV | Flaviviridae | *Dermacentor*<br>*Ixodes*<br>*Haemaphysalis* | 5,501 |

The top TBVs of concern represent seven virus families (*Asfarviridae*, *Flaviviridae*, *Nairoviridae*, *Orthomyxoviridae*, *Phenuiviridae*, *Poxviridae*, *Reoviridae*), two DNA viruses, and 12 RNA viruses (1 dsRNA, 11 ssRNA). Known tick vectors of these viruses include the genera *Amblyomma*, *Dermacentor*, *Haemaphysalis*, *Hyalomma*, *Ixodes*, *Ornithodoros*, *Rhipicephalus*. Human clinical symptoms from these viruses can be a mild, nonspecific febrile illness or can include severe, cognitive impairment^14,31^. Some of these viruses (Crimean-Congo hemorrhagic fever, Tick-borne encephalitis) are well documented, however many of these viruses have unknown enzootic cycles and limited knowledge of geographic risk^24,32^.

For our analyses, we restricted ZOVER records from the years 1990 to 2023 which reduced our observations to 3,845 and eliminated observations for Bhanja virus, Louping ill virus, Omsk hemorrhagic fever virus, and Sawgrass virus (Fig. 2). Tick-borne virus names were cleaned and filtered for our top 14 reported viruses of concern, referred to as tick-borne virus (TBV) hereafter (see Supplemental Text for more details on data cleaning). We began with the full country list from the package ‘rnaturaleath’ (n = 258 countries) and then excluded countries without reported square area in World Bank (indicator AG.LND.TOTL.K2) since most of them were small islands (n = 43)^40^. Each remaining country was assigned a binary code based on the presence (1) or absence (0) of a reported tick-virus in the ZOVER database (Supplemental Table 1). To address the additional bias introduced into the dataset, we obtain the number of citations per tick-borne virus species using the ‘easyPubMed’ package as a proxy for sampling effort^36,41^. We aggregate the citations for each tick-borne virus by country to capture the overall citation records and assess the relevance of each virus species (Supplemental Fig. 1).

**Figure 2.**
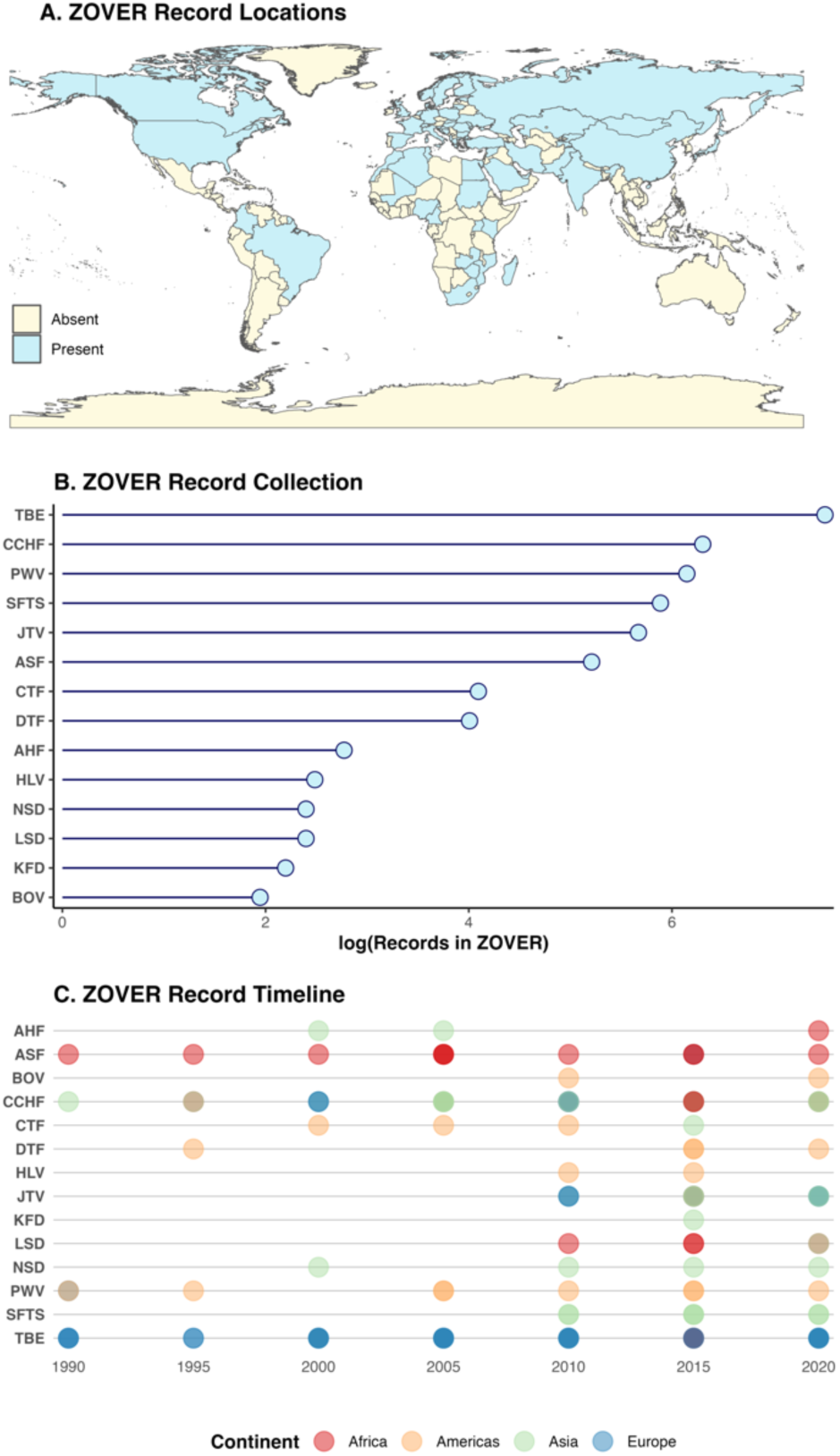
ZOVER database specifications used in this study: (**a**) Map of the top 14 reported tick-borne viruses for each country (record present-dark or absent-light), (**b**) total number of records for each tick-borne virus, and (**c**) timeline of reported tick-borne viruses recorded for every 5 years, color-coded by the continent of origin (Abbreviated tick-borne virus names given in Table 1).

#### Global trait matrix

For each country, when data was available (see Supplemental Table 2 for covariate coverage), we extracted a set of 23 covariates which fall under six broad categories: *environmental* suitability for ticks, *exposure* risk to ticks, likelihood of *knowledge* on tick-borne diseases, access to *health care*, *wealth* to attain care, and availability of *reporting* structures (see Table 2).

**Table 2.** Descriptions of covariates used for the BRT analyses.

| Driver Type | Variable | Description |
| --- | --- | --- |
| Environmental | Total number of ecoregions | Large areas defined by distinct natural communities, reflecting their original extent before significant land-use change |
| Environmental | Name of Köppen-Geiger Climate classification | Climate groups are defined by the types of vegetation in a specific region<sup>2,3</sup> |
| Environmental | Vegetation heterogeneity Index | Second-order dissimilarity of Enhanced Vegetation Index among neighboring pixels. Proxy for deforestation.<sup>4</sup> |
| Environmental | % of land forested | Forested area (% of total land area)<sup>5</sup> |
| Exposure | Livestock Production Index | An index measure of annual agricultural output relative to 2014–2016 base period<sup>6</sup> |
| Exposure | Livestock density | Gridded global distribution of cattle expressed in total number of cattle per pixel<sup>7</sup> |
| Exposure | % of population in agriculture | % of total employment involved in agriculture<sup>8</sup> |
| Knowledge | Pesticide imports | Current USD\$ spent on pesticide imports<sup>9</sup> |
| Knowledge | Presence of an International Veterinary Students Association (IVSA) Chapter | IVSA aims to benefit animals and humans globally by leveraging the skills and dedication of veterinary students to promote veterinary education and practice<sup>10</sup> |
| Knowledge | % of tertiary graduates from Ag, Vet, or Forestry schools | % of graduates from tertiary education graduating from Agriculture, Forestry, Fisheries and Veterinary programmes, both sexes (%)<sup>11</sup> |
| Knowledge | Adult Literacy Rate | % of people ages 15 and above literacy rate<sup>12</sup> |
| Knowledge | Education expenditures | % of GNI spent on education expenditures (adjusted savings)<sup>13</sup> |
| Health Care | Health care expenditures | % of GDP spent on healthcare expenditures<sup>14</sup> |
| Health Care | Reported average of annual people being treated for NTDs | The number of people needing treatment for neglected tropical diseases (NTDs) targeted by the WHO NTD Roadmap and reported to WHO, including those requiring mass treatment for PC-NTDs (lymphatic filariasis, onchocerciasis, schistosomiasis, soil-transmitted helminthiasis and trachoma).<sup>15</sup> |
| Wealth | Gini Index | Measure of income inequality within a population. Low values represent less inequality.<sup>16</sup> |
| Wealth | % of population living below national poverty line | Poverty headcount ratio at national poverty lines (% of population)<sup>17</sup> |
| Wealth | Social Vulnerability Index | A composite indicator, Subnational Human Development Index (SHDI), of relative multidimensional deprivation, based on a combination of three dimensions: education, health, and standard of living. Low values represent less vulnerability.<sup>18</sup> |
| Wealth | % of population with access to electricity | % of population with access to electricity<sup>19</sup> |
| Reporting | Human population counts | Gridded population counts<sup>20</sup> |
| Reporting | Land area (sq km) | A country's total area, excluding inland water bodies, continental shelf claims, and exclusive economic zones.<sup>21</sup> |
| Reporting | Conflict exposure (total events) | Population that is living in an area of active disorder, where people may be directly injured, caught in conflict, targeted, or affected by the destruction of their community.<sup>22</sup> |
| Reporting | Conflict exposure (% of population exposed) | ‘Conflict exposure’ is a measure of the number of people living within 1 kilometer, 2 km, and 5 km of each conflict incident or demonstration.<sup>22</sup> |
| Reporting | % of population in urban areas | % of total population living in urban areas<sup>23</sup> |

Covariates included the mode Köppen-Geiger Climate Classification within a country^42,43^, total number of ecoregions within a country^44^, mean vegetation heterogeneity per country^45^, % of total land area that is forested (World Bank indicator AG.LND.FRST.ZS) (environment category); annual aggregate volume of livestock production (World Bank indicator AG.PRD.LVSK.XD), livestock density^46^, % of population employed in agriculture (WorldBank indicator SL.AGR.EMPL.ZS), (exposure category); annual pesticide imports (USD$) (WorldBank indicator BM.AG.PEST.CD), presence of an International Veterinary Students Association chapter (IVSA) (https://www.ivsa.org), % of graduates from tertiary education graduating from Agriculture, Forestry, Fisheries, and Veterinary programs (WorldBank indicator SE.TER.GRAD.AG.ZS), adult literacy rate (WorldBank indicator SE.ADT.LITR.ZS), education expenditures (% of GNI) (WorldBank indicator NY.ADJ.AEDU.GN.ZS) (knowledge category); health care expenditures (% of GDP) (WorldBank indicator SH.XPD.CHEX.GD.ZS), reported average of annual people being treated for neglected tropical diseases (World Health Organization indicator SDGNTDTREATMENT) (health care category); social vulnerability index (CIESIN GRDI), % of population living below national poverty line (WorldBank indicator SI.POV.NAHC), % of population with access to electricity (WorldBank indicator EG.ELC.ACCS.ZS), Gini Index (WorldBank indicator SI.POV.GINI) (wealth category); conflict exposure to total events^47^, % of population exposed to conflict^47^, human population count (CIESIN Gridded Population), total area of country (WorldBank indicator AG.LND.TOTL.K2), % of population living in urban areas (WorldBank indicator SP.URB.TOTL.IN.ZS) (reporting category) (Table 2, Supplemental Table 3). When possible, covariate data was averaged between 2010-2020 for each country, or the most recent year available was chosen. See Supplemental Text for further details on data decisions.

### Statistical Analysis

We applied generalized boosted regression trees (BRT), a machine learning algorithm, to classify country-level reported TBV outcomes. With this method and our global trait matrix we can identify environmental and socioeconomic characteristics of countries with reported TBVs compared to countries that did not have reported TBVs. Boosted regression trees can handle non-normal, highly collinear, and non-randomly missing covariates better than traditional regression methods^35–37^. This approach identifies patterns that could generate more formal predictions to guide future research on potential socio-ecological levers to improve diagnostics and reporting of tick-borne viruses^35–37,48^.

We used BRTs with Bernoulli-distristributed error for our binary response. For our predictors, we removed covariates that had low coverage (< 50%) and minimal variation (> 97% homogenous)^36,48^. For covariates that were highly skewed (i.e., total area of country, human population counts), we took the log_10_ transformation value. Before analysis, we randomly divided the data into training (70%) and test (30%) sets using the ‘rsamplè package to ensure stratified sampling in equal proportions^48,49^. To determine the optimal values for model parameters such as number of trees, learning rate, and interaction depth and to avoid potential overfitting, a grid search was conducted for hyperparameter tuning^36,48^. We fit the BRT models using the ‘gbm’ package and applied five-fold cross validation^50^. To control for the potential effects of sampling bias on our results, we use the total citation number from ‘easyPubMed’ package as a Poisson distributed outcome for a second set of BRT models using the same hyperparameters^36,37,48^.

To determine model performance we used the ‘rocr’ package to derive model metrics such as accuracy as the area under the receiver operator curve (AUC)^51^ and ‘InformationValuè package for sensitivity and specificity^52^. Since the results may vary based on random splits between training and test data, we employed 100 stratified partitions to create an ensemble yielding mean performance measures (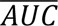 for accuracy; 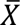 for specificity and sensitivity)^48^. To identify the influence of each predictor on our outcome – country-level reported tick-borne virus – we look at the relative variable importance, which indicates the proportion of improvement in model fit that is attributed to each predictor^36^. To visualize the marginal effects of an individual predictor variable on our outcome, while controlling for all other predictors, we created partial dependence plots^36^.

## Results

### Dataset description

A total of 59 countries have a reported TBV of top concern from the ZOVER database (Fig 2, Supplemental Table 1). The country with the most reported unique TBVs was China (n = 6), followed by the United States (n = 5), and countries with at least three unique TBVs included Egypt, France, India, Japan, Kazakhstan, Kenya, Russia, and Serbia. However, the countries with the most recorded TBV observations were Russia (n = 1,466), the United States (n = 580), and China (n = 278). The most reported TBVs were Tick-borne encephalitis (n = 1,818), Crimean-Congo hemorrhagic fever (n = 546), and Powassan virus (n = 467). Most countries in our dataset only had 1 unique TBV (n = 45).

### Trait profile of countries with reported tick-borne viruses

To identify characteristics associated with tick-borne virus reported outcomes, we evaluated the relative importance of 23 variables with boosted regression tree (BRT) models. Our BRT model distinguished reported vs unreported cases with relatively high accuracy (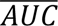= 0.87 ± 0.0038) and specificity 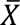 = 0.90 ± 0.0045), but low sensitivity 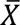 = 0.49 ± 0.013). The mean relative influence of all variables was 4.17 ± 29. The top three influential variables were Köeppen-Geiger climate classification zone (relative influence = 24%), Gini Index (relative influence = 11%), and human population counts (relative influence = 9%). Other relatively important variables included pesticide imports (USD$), land area (sq km), livestock production index, and conflict exposure to total events (Fig. 3, Supplemental Table 3). Partial dependence plots (Fig. 4, Supplemental Fig. 2) described countries that were more likely to report a TBV had a lower Gini Index (i.e., countries with less inequalities such as Nordic countries), increased dollars spent on pesticide imports, and had institutions (i.e., IVSA chapter) or individuals with agricultural, forestry, or veterinary knowledge (i.e., % of tertiary grads) present. Additional characteristics included countries with a lower percent of population exposed to conflict also had a higher probability of reporting a TBV. None of our variables had a relative influence of 0%, however the percentage of population with access to electricity had a relative influence of 0.42% (Supplemental Table 4). For our BRT models with a Bernoulli response (i.e., reported TBV), we identified an optimal learning rate of 0.01, interaction depth of 4, and a maximum number of trees of 5,000 (Supplemental Fig. 3). When we ran a separate BRT model with citation weight per country as a proxy for sampling effort, we found that citation weight was not predicatable by our global trait matrix (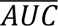 = 0.50 ± 4.1 *x* 10^−7^) suggesting that the covariates associated with reported TBVs were not confounded by the traits of well-studied viruses within a given country. For our BRT model with a Poisson response (i.e., citation weight per country), we identified an optimal learning rate of 0.0005, interaction depth of 2, and a maximum number of trees of 15,000.

**Figure 3.**
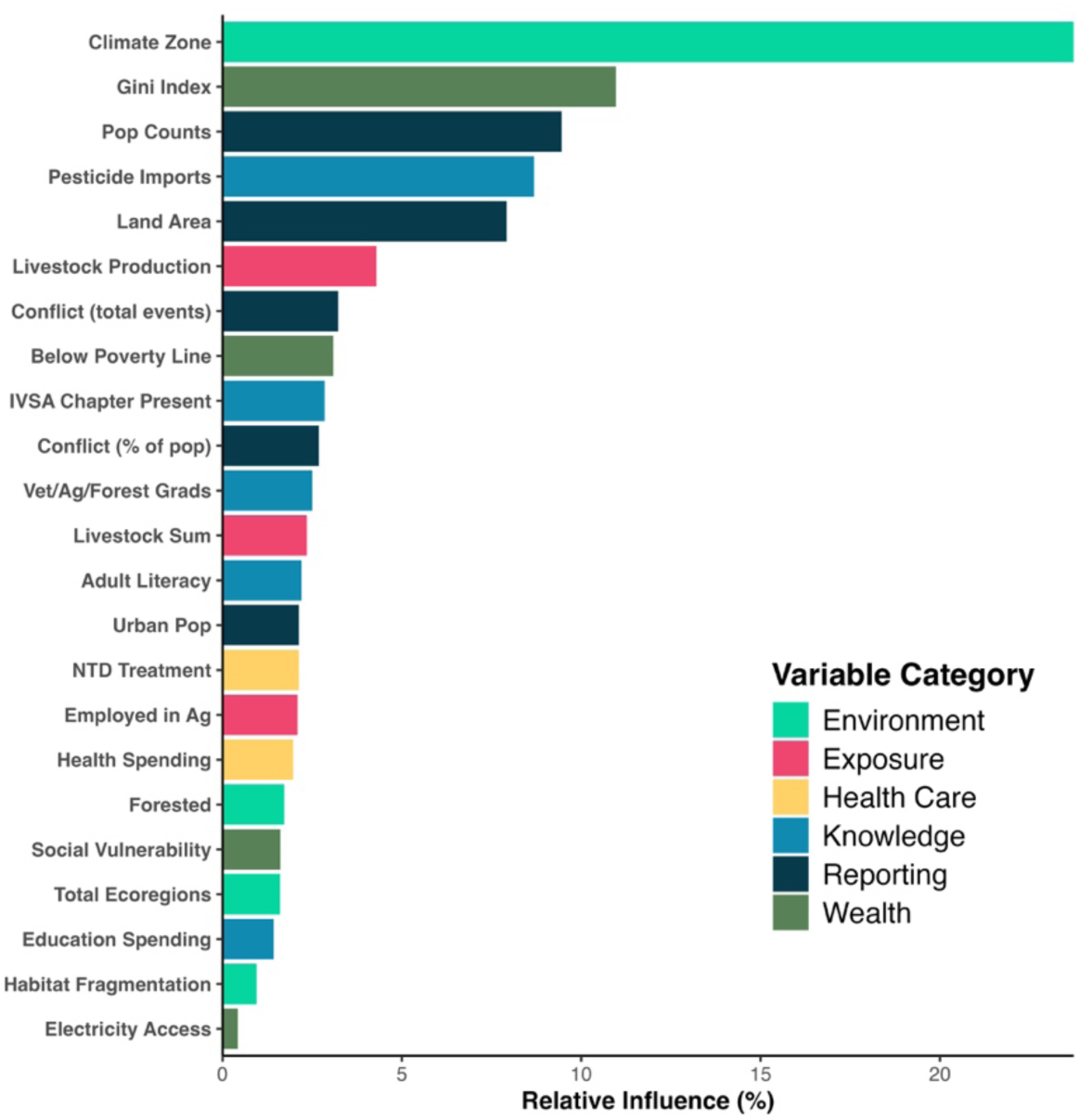
Variable importance values for the covariates in the BRT model. The value represents the proportion of improvement in model fit that is attributed to each predictor.

**Figure 4.**
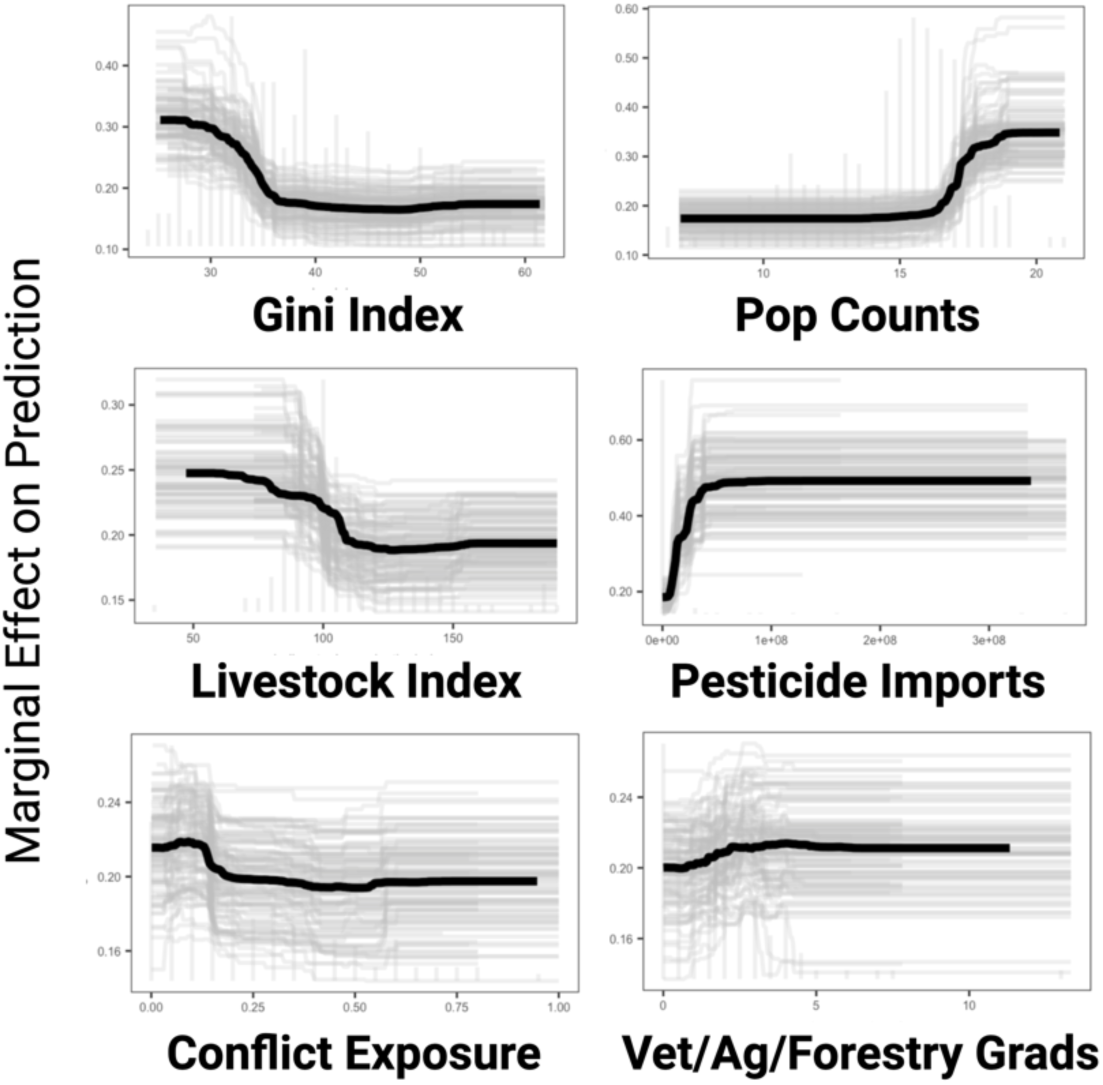
Partial dependence plots of some of the top BRT numeric predictors (Gini Index, human population counts, livestock production index, pesticide imports, % of population exposed to conflict, % of tertiary grads from veterinary, agricultural, or forestry graduate programs). Histograms show the distribution of the underlying covariates. The black line displays the marginal effect of a given variable while controlling for all other predictors for predicting a country with a reported tick-virus.

## Discussion

Public health resources and attention is allocated based on the relative magnitude of a number, specifically reported disease cases. However, reported incidence may not reflect true disease risk. Disease risk is a combination of environmental suitability for a pathogen to propagate and transmit as well as socioeconomic factors that influence the contact rates between pathogens and humans. Our study highlights the importance of both environmental and socioeconomic drivers of reported tick-borne viruses. Similar to other infectious diseases, both types of drivers are important to the transmission of infectious diseases. However, our study highlights the inequities in reporting infectious diseases, specifically tick-borne viruses, that require additional resources such as sufficient knowledge, adequate testing, and wealth to attain care. Due to the results of our study, we suggest socioeconomic interventions to be applied to countries with suspected TBVs to reduce the burden of tick-borne viruses globally. In accordance with other infectious diseases, by improving societal conditions, you can improve the health of all people.

Environmentally mediated pathogens, such as tick-borne viruses that are vectored by free-living ectotherms, pose a significant global health burden^8^. However, unlike directly transmitted diseases (e.g., influenza, tuberculosis, measles), environmentally mediated pathogens allow for more interventions that can disrupt transmission to humans before requiring conventional medical approaches^4^. We find that countries are more likely to report a TBV if they were wealthy and had institutions (i.e., IVSA chapter) or individuals with agricultural, forestry, or veterinary knowledge (i.e., % of tertiary grads) present. Additional characteristics included countries with a lower percent of population exposed to conflict also had a higher probability of reporting a TBV. For a group of diseases that require specific knowledge and readily available diagnostic tools, our results make intuitive sense on a country-level^4,5,27^. If a country is unable to meet the basic needs of its citizens (i.e., high social vulnerability index, high percentage of population exposed to conflict, high percentage of population lives below the national poverty line) then it is unlikely to have resources for standardized surveillance to recognize and treat diseases that have less public health campaign efforts and less defined clinical symptoms^5,7,24^.

Although much literature has supported that global disease burdens are concentrated in communities with less economic resources, environmental burden may not translate to reported disease cases^5,19,20^. Our work is supported by other country-level analysis of socio-economic-ecological drivers of environmentally mediated disease which found more affluent communities were more likely to report disease cases^4,5,7^. Potential mechanisms of this relationship include people in wealthier nations living in cities which may reduce distance to medical facilities and allow for a more efficient public health campaign^4,5,16,18^.

Although country-level analyses are more likely to match the scale most relevant for policy making, rather than sub-country level, it has its limitations^4,5^. Country-level analyses obscures climate patterns, which are indisputably important components for the distribution and seasonality of exposure to TBVs^24^. We tried to include environmental covariates that would represent ecological conditions possible for a tick to be present such as Köppen-Geiger Climate Classification zones, which our model did identify as a relatively important variable, and specifically humid subtropical climate, humid continental climate, and hot summer Mediterranean climate had higher probability of reporting a tick-virus. Our environmental results are supported by a recent study that also identified land cover identity as a superior method for predicting areas suitable for ticks compared to climate data alone^53^. However, some of these relationships may have been skewed because we took the mode classification zone per country (e.g., subarctic climate for Russia, Finland, Sweden) (Supplemental Fig. 3). Our knowledge from this study is also as only good as the knowledge contributed to the ZOVER database, which inherently introduces biases and part of the problem of interpreting reported cases for tick-borne diseases, as well as covariate coverage (Supplemental Fig. 4). Attempts to reduce the magnitude of bias were undertaken via modeling our outcome as a binary presence/absence record.

Additional steps taken were the inclusion of an additional outcome of citation weight per country (number of easyPubMed citations per virus totaled per country) and focusing on TBVs with well-established etiology and transmission pathways (Supplemental Fig. 1). Our model with the citation weight outcome was not predicted (i.e., test AUC = 50) by the same set of covariates used in the BRT model with the binary reported TBV, suggesting that our results from our primary model were not confounded by sampling biases. Notable countries absent from the ZOVER database prior to data quality filtering included Mexico, Tanzania, and many central American countries with known tick-viruses. This points to a need for further efforts to document and collect additional data for unique and singular occurrence data on ZOVER as well as expand surveillance in geographic areas that are presumed to be ecologically suitable for TBVs to occur^33,53^.

Tick-borne diseases are a function of a complex chain of events, spanning the remit of multiple fields of investigation and understanding. Thus, they are suited to One Health framed approaches to best capture their dynamics and occurrence, and to thereby inform mitigation or intervention^24,32,33^. Ticks and the pathogens they harbor can thrive in various environments, including natural communities (such as enzootic cycles without human hosts), peri-urban areas (where humans and companion animals coexist), and economic activities (like animal husbandry, agriculture, and forestry). Due to this wide network of exposure, physicians, veterinarians, and occupational health experts should work together to develop, maintain, and share tick surveillance to achieve the common goal of reducing human disease burden^32^. A One Health surveillance system could strengthen communication between these groups to improve tick identification, distribution of diagnostic tests, and implementation of tick-prevention strategies^28,54^. While the creation of a database like ZOVER is a positive step toward reducing tick-borne diseases, it is crucial to enhance our understanding of the real environmental risks posed by TBVs through robust active surveillance of ticks and their natural hosts.

### Conclusion

Launched in 2022 by the World Health Organization, the Global Arbovirus Initiative aims to address the increasing risk of arbovirus epidemics, including tick-borne viruses. The initiative focuses on monitoring disease risk, strengthening vector control, and building a coalition of partners. Our study identifies potential socio-ecological levers to improve the reporting of tick-borne viruses at a country level. We find countries that were more likely to report TBVs had a lower Gini Index (i.e., countries with less inequalities such as Nordic countries), increased dollars spent on pesticide imports, and had institutions (i.e., IVSA chapter) or individuals with agricultural, forestry, or veterinary knowledge (i.e., % of tertiary grads) present. Additional characteristics included countries with a lower percent of population exposed to conflict also had a higher probability of reporting a TBV. Although the environmental suitability for a pathogen to persist is important, our study suggests policy aimed at economic development may be the most effective tool to reduce tick-borne viruses.

## Acknowledgements

SS and SJR were supported by funding to the Viral Emergence Research Initiative (VERENA) Institute, including NSF BII 2021909 & NSF BII 2213854, as part of a Fellow-In-Residence program.

## Data Sharing

ZOVER data is freely available under the CC BY-NC license version 4.0. Code on data processing, BRTs, and visualizations can be found on https://github.com/sbsambado/global_tbvs.

## Data Availability

Sources of data used in this study are publicly available and cited in the manuscript text, Suppelemental Table 5. Additional citation information: (**WorldBank**) Specific World development indicators are listed in Method Section https://databankfiles.worldbank.org/public/ddpext/; (**CIESIN GRDI**) Center for International Earth Science Information Network - CIESIN - Columbia University. 2022. Global Gridded Relative Deprivation Index, Version 1. Palisades, New York: NASA Socioeconomic Data and Applications Center (SEDAC). https://doi.org/10.7927/3xxe-ap97<u>. Accessed 28 May 2024</u>; (**CIESIN Gridded Population**) Center for International Earth Science Information Network - CIESIN - Columbia University. 2016. Gridded Population of the World, Version 4 (GPWv4): Population Count. Palisades, NY: NASA Socioeconomic Data and Applications Center (SEDAC). http://dx.doi.org/10.7927/H4X63JVC. Accessed 02 June 2024; (**Conflict Exposure**) Armed Conflict Location and Event Data Project (ACLED and WorldPop) 27 September 2024. Conflict Exposure to Political Violence (Battles, Explosions/Remote violence, Violence against civilians, Excessive force against protesters, & Mob violence). Best gridded estimate (1 km - 5 km); (**NCBI Taxonomy Browser**) https://www.ncbi.nlm.nih.gov/Taxonomy/Browser/ www.tax.cgi.

## Article Notes

The authors declare that there are no competing financial interests.

