## Supplemental Material for "Ecological and socioeconomic factors associated with globally reported tick-borne viruses"

Table of Contents

**S Text**. Further details on data processing and model decisions

**S Figure 1**. Bar graph of citation weight per country

**S Figure 2**. Additional partial dependence plots

**S Figure 3**. Performance of parameters in the hypertuning grid

**S Figure 4**. Global distribution of important covariates

**S Table 1**. List of countries included in analysis

**S Table 2**. Coverage of covariates included in models

**S Table 3**. Additional covariate source information

**S Table 4.** Variable importance rank

**S Text**.

*Tick-virus occurrence data*

Tick-virus data came from the most comprehensive current database on tick-borne viruses (ZOVER; <http://www.mgc.ac.cn/cgi-bin/ZOVER/main.cgi>). On ZOVER, we went to the page for tick-associated viruses (<http://www.mgc.ac.cn/cgi-bin/ZOVER/mainTable.cgi?db=tick>).  On the tick-associated viruses page, under ‘Browse by virus’, for each virus grouping (i.e., ssDNA viruses, dsRNA viruses, dsDNA viruses no RNA stage, unclassified viruses, ssRNA negative-strand viruses, and ssRNA positive-strand viruses, no DNA stage) we click Show ‘All’ entries, and ‘Save this table’ as xls files. Data was accessed on 2024-09-21. Data cleaning steps can be found in ‘2_ZOVEROutcomes _RawtoClean.Rmd’ on GitHub repository <https://github.com/sbsambado/global_tbvs>. Specifically, for the collection year, if the year was reported as Year 1 ~ Year 2, we chose the first year (Year 1). We then aggregate observations by five years, with the exception of observations before 1950. Further steps were taken to ensure correct tick virus names for the top 18 viruses using a series of string commands in RStudio (e.g., grepl(), str_detect(), case_when()). To filter for our top tick-borne viruses of concern the following strings were used to filter for observations in ZOVER: “African swine”, “Alkhumra”, “Bhanja”, “Bourbon”, “Colorado”, “Crimean”, “Deer tick”, “Heart”, “Jingmen”, “Kyasanur”, “Louping”, “Lumpy”, “Nairobi”, “Omsk”, “Powassan”, “Saw”, “Severe”, and “Tick-borne enceph”.

*Global trait matrix data*

To ensure that multiple data sources could easily be merged, much attention was given to country names for each data source. We based our final country names on the package ‘rnaturalearth’ using the command ne_countries(scale = “large”, type = “countries”, returnclass = “sf”). This resulted in 258 countries including US Naval Base Guantanamo Bay, Southern Patagonian Ice Field, and other countries that were mainly small islands. However, all covariate sources were matched against the ‘rnaturalearth’ country list and if there was no observation for a country an NA was assigned for that particular covariate. Data cleaning and processing steps can be found in ‘1_TraitMatrix_RawtoClean.Rmd’ on GitHub repository <https://github.com/sbsambado/global_tbvs>. Specifically, data from World Bank (e.g., land area, Gini Index, adult literacy rate, below national poverty line, health expenditure) were selected from 2018 or the most recent year of observation. For the adult literacy rate, many countries were not reported but some assumptions were made about G20 countries having a 99.9% adult literacy rate to ensure this covariate could be included in boosted regression tree analysis. For covariate data that was derived from gridded remotely sensed data, data was extracted from ‘rnaturalearth’ country polygons.

**S Figure 1**. Citation weight assigned to each country with reported tick-virus broken down by (A) country and (B) tick-virus. PubMed and Citation weights were log transformed.

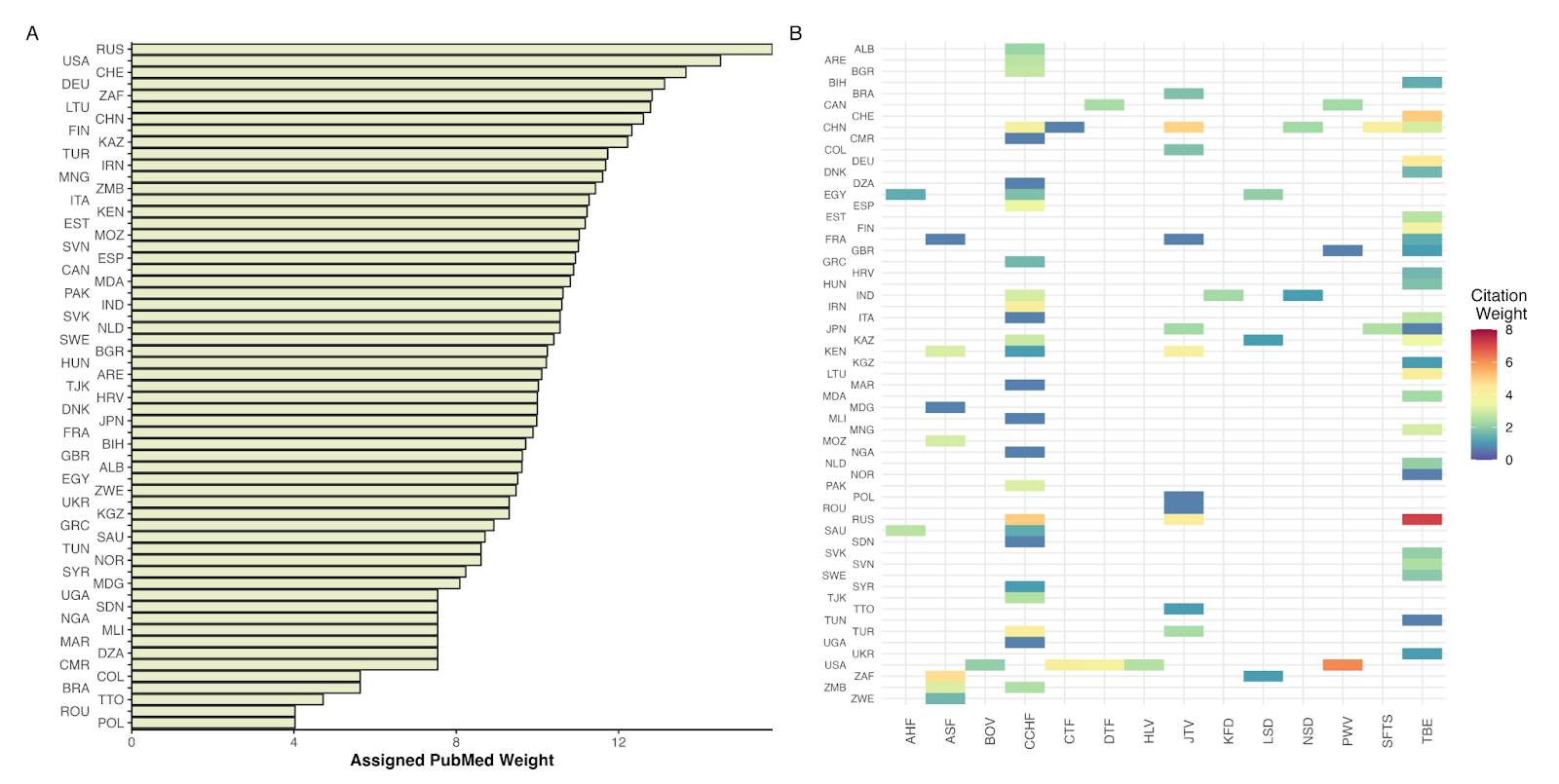

**S Figure 2.** Additional partial dependence plots of the response variable on the predictor (1 = tick-virus present in a country), indicating how the predicted values changes as the predictor variable increases. The short verticle ticks along the x-axis indicate the distribution of the numeric predictor variables grouped by deciles. The highest values for Köeppen-Geiger Climate Classification Zones include subartic climate (Dfc), marine climate (Cfc), humid subtropical climate (Cfa), and humid continental climate (Dwb).

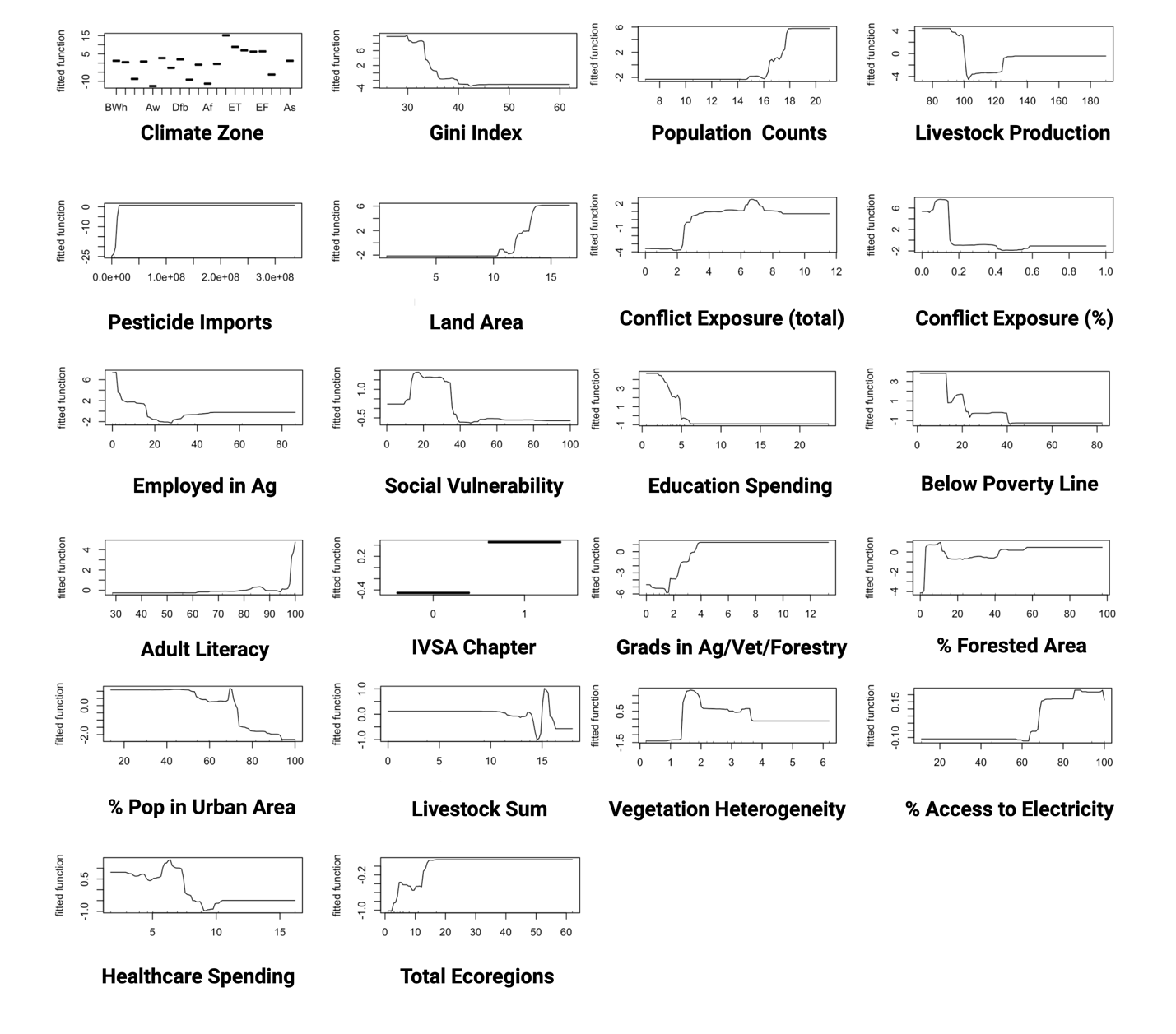

**S Figure 3.** Performance of parameters in the hyperparameter tuning grid for BRT models. The plot displays test AUC by learning rate, with colors representing interaction depth (number of trees = 5,000).

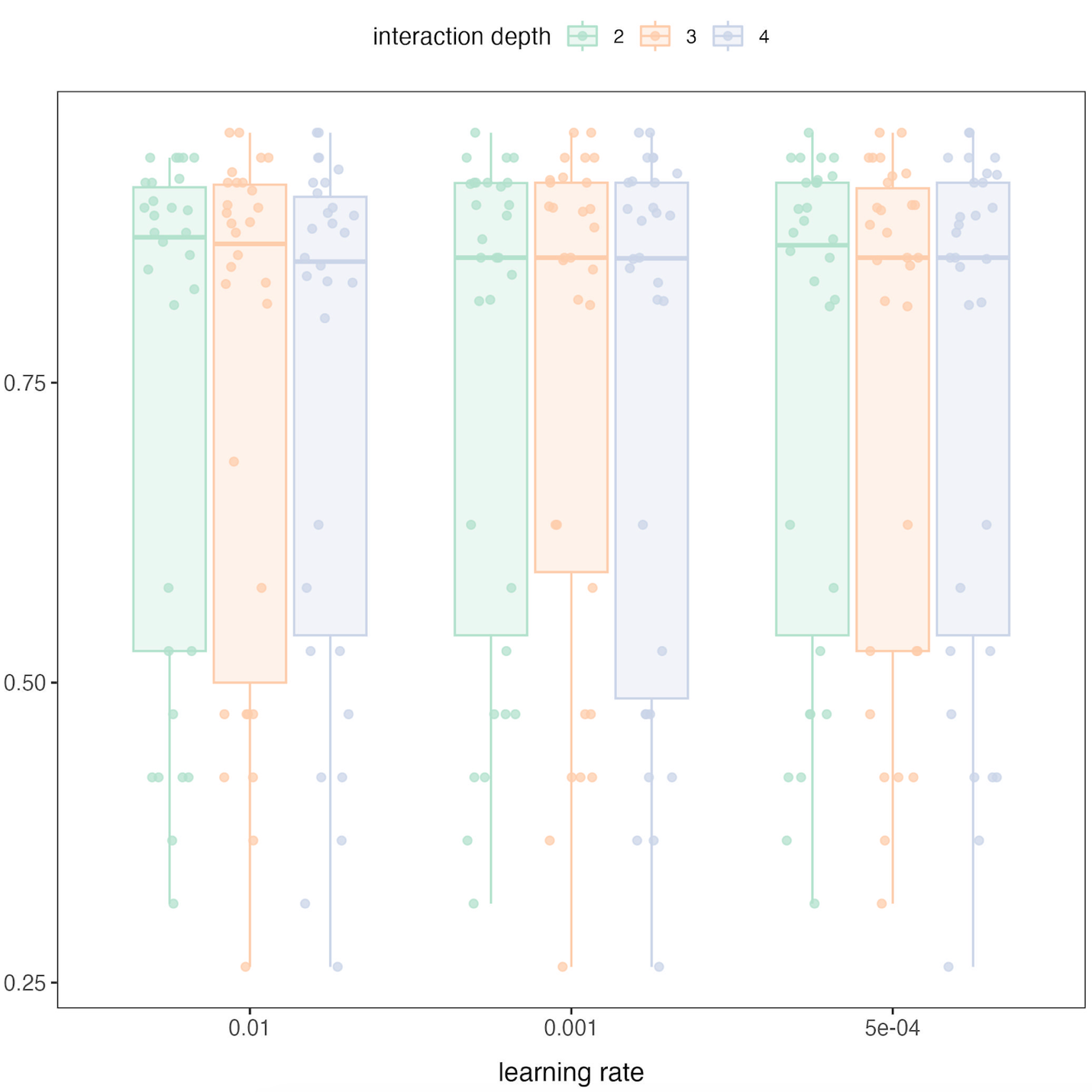

**S Figure 4.** Global distribution of relatively important variables, where colors indicate the intensity of the reported variable (dark = high values, light = low values, beige = NA values).

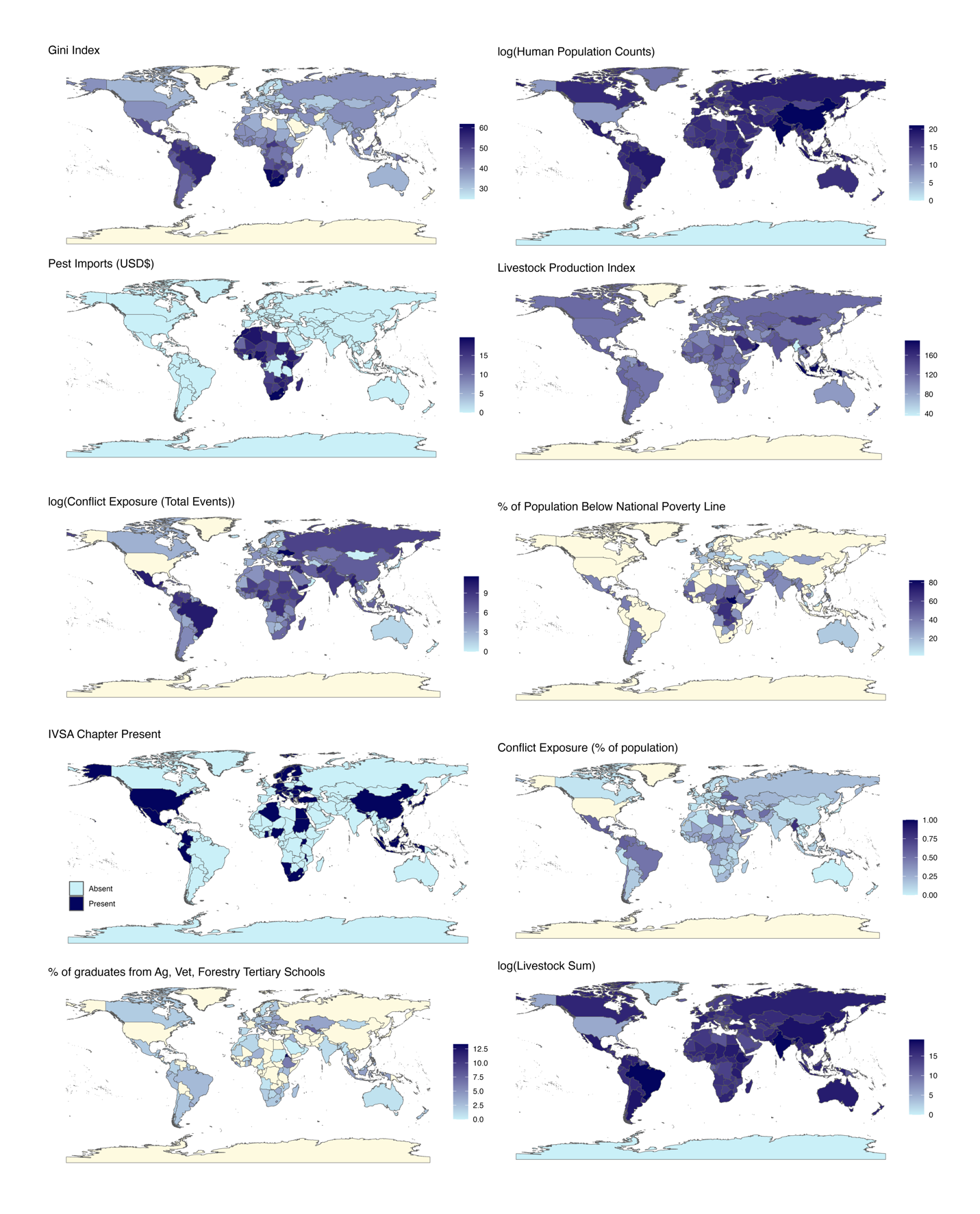

**S Figure 5**. Map of ZOVER coverage for (A) selected tick-borne viruses (TBVs) from 1990-2023, and (B) all TBVs between 1950-2023, where colors indicate the intensity of the reported variable (dark represents high values, light indicates low values, and beige denotes countries with NA values)

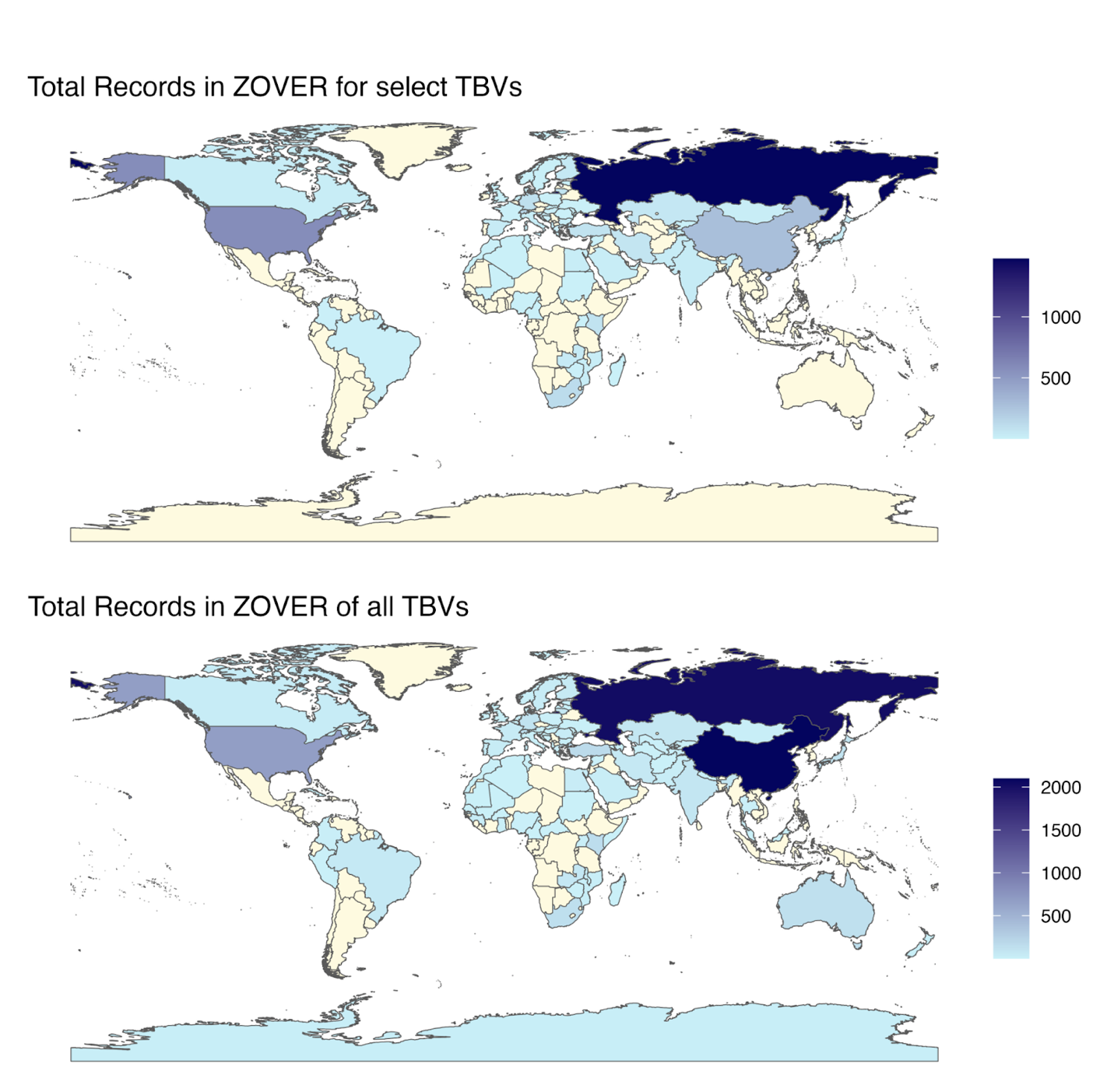

**S Table 1**.  List of TBVs by country. Tick-virus abbreviations: CCHF = Crimean-Congo hemorrhagic fever, TBE = Tick-borne encephalitis, JTV = Jingmen tick virus, DTF = Deer tick virus, PWV = Powassan virus, CTF = Colorado tick fever virus, NSD = Nairobi sheep disease virus, SFTS = Severe fever with thrombocytopenia syndrome virus, AHF = Alkhumara hemorrhagic fever virus, LSD = Lumpy skin disease virus, ASF = African swine fever virus, BHV = Bourbon virus, KFD = Kyasanur forest disease virus,  HLV = Heartland virus.

| **Admin** | **ISO A3** | **CCHF** | **TBE** | **JTV** | **DTF** | **PWV** | **CTF** | **NSD** | **SFTS** | **AHF** | **LSD** | **ASF** | **KFD** | **BOV** | **HLV** |
| --- | --- | --- | --- | --- | --- | --- | --- | --- | --- | --- | --- | --- | --- | --- | --- |
| **Albania** | ALB | 8 | 0 | 0 | 0 | 0 | 0 | 0 | 0 | 0 | 0 | 0 | 0 | 0 | 0 |
| **Algeria** | DZA | 1 | 0 | 0 | 0 | 0 | 0 | 0 | 0 | 0 | 0 | 0 | 0 | 0 | 0 |
| **Bosnia and Herzegovina** | BIH | 0 | 3 | 0 | 0 | 0 | 0 | 0 | 0 | 0 | 0 | 0 | 0 | 0 | 0 |
| **Brazil** | BRA | 0 | 0 | 5 | 0 | 0 | 0 | 0 | 0 | 0 | 0 | 0 | 0 | 0 | 0 |
| **Bulgaria** | BGR | 15 | 0 | 0 | 0 | 0 | 0 | 0 | 0 | 0 | 0 | 0 | 0 | 0 | 0 |
| **Cameroon** | CMR | 1 | 0 | 0 | 0 | 0 | 0 | 0 | 0 | 0 | 0 | 0 | 0 | 0 | 0 |
| **Canada** | CAN | 0 | 0 | 0 | 10 | 9 | 0 | 0 | 0 | 0 | 0 | 0 | 0 | 0 | 0 |
| **China** | CHN | 52 | 20 | 138 | 0 | 0 | 1 | 9 | 58 | 0 | 0 | 0 | 0 | 0 | 0 |
| **Colombia** | COL | 0 | 0 | 5 | 0 | 0 | 0 | 0 | 0 | 0 | 0 | 0 | 0 | 0 | 0 |
| **Croatia** | HRV | 0 | 4 | 0 | 0 | 0 | 0 | 0 | 0 | 0 | 0 | 0 | 0 | 0 | 0 |
| **Czech Republic** | NA | 0 | 18 | 0 | 0 | 0 | 0 | 0 | 0 | 0 | 0 | 0 | 0 | 0 | 0 |
| **Denmark** | DNK | 0 | 4 | 0 | 0 | 0 | 0 | 0 | 0 | 0 | 0 | 0 | 0 | 0 | 0 |
| **Egypt** | EGY | 5 | 0 | 0 | 0 | 0 | 0 | 0 | 0 | 3 | 7 | 0 | 0 | 0 | 0 |
| **Estonia** | EST | 0 | 13 | 0 | 0 | 0 | 0 | 0 | 0 | 0 | 0 | 0 | 0 | 0 | 0 |
| **Finland** | FIN | 0 | 41 | 0 | 0 | 0 | 0 | 0 | 0 | 0 | 0 | 0 | 0 | 0 | 0 |
| **France** | France | 0 | 3 | 1 | 0 | 0 | 0 | 0 | 0 | 0 | 0 | 1 | 0 | 0 | 0 |
| **Germany** | DEU | 0 | 92 | 0 | 0 | 0 | 0 | 0 | 0 | 0 | 0 | 0 | 0 | 0 | 0 |
| **Greece** | GRC | 4 | 0 | 0 | 0 | 0 | 0 | 0 | 0 | 0 | 0 | 0 | 0 | 0 | 0 |
| **Hungary** | HUN | 0 | 5 | 0 | 0 | 0 | 0 | 0 | 0 | 0 | 0 | 0 | 0 | 0 | 0 |
| **India** | IND | 20 | 0 | 0 | 0 | 0 | 0 | 2 | 0 | 0 | 0 | 0 | 9 | 0 | 0 |
| **Iran** | IRN | 63 | 0 | 0 | 0 | 0 | 0 | 0 | 0 | 0 | 0 | 0 | 0 | 0 | 0 |
| **Italy** | ITA | 1 | 14 | 0 | 0 | 0 | 0 | 0 | 0 | 0 | 0 | 0 | 0 | 0 | 0 |
| **Japan** | JPN | 0 | 1 | 9 | 0 | 0 | 0 | 0 | 11 | 0 | 0 | 0 | 0 | 0 | 0 |
| **Kazakhstan** | KAZ | 17 | 31 | 0 | 0 | 0 | 0 | 0 | 0 | 0 | 2 | 0 | 0 | 0 | 0 |
| **Kenya** | KEN | 2 | 0 | 50 | 0 | 0 | 0 | 0 | 0 | 0 | 0 | 21 | 0 | 0 | 0 |
| **Korea, South** | NA | 0 | 20 | 0 | 0 | 0 | 0 | 0 | 291 | 0 | 0 | 0 | 0 | 0 | 0 |
| **Kyrgyzstan** | KGZ | 0 | 2 | 0 | 0 | 0 | 0 | 0 | 0 | 0 | 0 | 0 | 0 | 0 | 0 |
| **Lithuania** | LTU | 0 | 65 | 0 | 0 | 0 | 0 | 0 | 0 | 0 | 0 | 0 | 0 | 0 | 0 |
| **Madagascar** | MDG | 0 | 0 | 0 | 0 | 0 | 0 | 0 | 0 | 0 | 0 | 1 | 0 | 0 | 0 |
| **Mali** | MLI | 1 | 0 | 0 | 0 | 0 | 0 | 0 | 0 | 0 | 0 | 0 | 0 | 0 | 0 |
| **Moldova** | MDA | 0 | 9 | 0 | 0 | 0 | 0 | 0 | 0 | 0 | 0 | 0 | 0 | 0 | 0 |
| **Mongolia** | MNG | 0 | 20 | 0 | 0 | 0 | 0 | 0 | 0 | 0 | 0 | 0 | 0 | 0 | 0 |
| **Morocco** | MAR | 1 | 0 | 0 | 0 | 0 | 0 | 0 | 0 | 0 | 0 | 0 | 0 | 0 | 0 |
| **Mozambique** | MOZ | 0 | 0 | 0 | 0 | 0 | 0 | 0 | 0 | 0 | 0 | 19 | 0 | 0 | 0 |
| **Netherlands** | NLD | 0 | 7 | 0 | 0 | 0 | 0 | 0 | 0 | 0 | 0 | 0 | 0 | 0 | 0 |
| **Nigeria** | NGA | 1 | 0 | 0 | 0 | 0 | 0 | 0 | 0 | 0 | 0 | 0 | 0 | 0 | 0 |
| **Norway** | Norway | 0 | 1 | 0 | 0 | 0 | 0 | 0 | 0 | 0 | 0 | 0 | 0 | 0 | 0 |
| **Pakistan** | PAK | 22 | 0 | 0 | 0 | 0 | 0 | 0 | 0 | 0 | 0 | 0 | 0 | 0 | 0 |
| **Poland** | POL | 0 | 0 | 1 | 0 | 0 | 0 | 0 | 0 | 0 | 0 | 0 | 0 | 0 | 0 |
| **Romania** | ROU | 0 | 0 | 1 | 0 | 0 | 0 | 0 | 0 | 0 | 0 | 0 | 0 | 0 | 0 |
| **Russia** | RUS | 152 | 1259 | 66 | 0 | 0 | 0 | 0 | 0 | 0 | 0 | 0 | 0 | 0 | 0 |
| **Saudi Arabia** | SAU | 3 | 0 | 0 | 0 | 0 | 0 | 0 | 0 | 13 | 0 | 0 | 0 | 0 | 0 |
| **Serbia** | NA | 41 | 1 | 2 | 0 | 0 | 0 | 0 | 0 | 0 | 0 | 0 | 0 | 0 | 0 |
| **Slovakia** | SVK | 0 | 7 | 0 | 0 | 0 | 0 | 0 | 0 | 0 | 0 | 0 | 0 | 0 | 0 |
| **Slovenia** | SVN | 0 | 11 | 0 | 0 | 0 | 0 | 0 | 0 | 0 | 0 | 0 | 0 | 0 | 0 |
| **South Africa** | ZAF | 0 | 0 | 0 | 0 | 0 | 0 | 0 | 0 | 0 | 2 | 114 | 0 | 0 | 0 |
| **Spain** | ESP | 30 | 0 | 0 | 0 | 0 | 0 | 0 | 0 | 0 | 0 | 0 | 0 | 0 | 0 |
| **Sudan** | SDN | 1 | 0 | 0 | 0 | 0 | 0 | 0 | 0 | 0 | 0 | 0 | 0 | 0 | 0 |
| **Sweden** | SWE | 0 | 6 | 0 | 0 | 0 | 0 | 0 | 0 | 0 | 0 | 0 | 0 | 0 | 0 |
| **Switzerland** | CHE | 0 | 156 | 0 | 0 | 0 | 0 | 0 | 0 | 0 | 0 | 0 | 0 | 0 | 0 |
| **Syria** | SYR | 2 | 0 | 0 | 0 | 0 | 0 | 0 | 0 | 0 | 0 | 0 | 0 | 0 | 0 |
| **Tajikistan** | TJK | 12 | 0 | 0 | 0 | 0 | 0 | 0 | 0 | 0 | 0 | 0 | 0 | 0 | 0 |
| **Trinidad and Tobago** | TTO | 0 | 0 | 2 | 0 | 0 | 0 | 0 | 0 | 0 | 0 | 0 | 0 | 0 | 0 |
| **Tunisia** | TUN | 0 | 1 | 0 | 0 | 0 | 0 | 0 | 0 | 0 | 0 | 0 | 0 | 0 | 0 |
| **Turkey** | TUR | 66 | 0 | 10 | 0 | 0 | 0 | 0 | 0 | 0 | 0 | 0 | 0 | 0 | 0 |
| **Uganda** | UGA | 1 | 0 | 0 | 0 | 0 | 0 | 0 | 0 | 0 | 0 | 0 | 0 | 0 | 0 |
| **Ukraine** | UKR | 0 | 2 | 0 | 0 | 0 | 0 | 0 | 0 | 0 | 0 | 0 | 0 | 0 | 0 |
| **United Arab Emirates** | ARE | 13 | 0 | 0 | 0 | 0 | 0 | 0 | 0 | 0 | 0 | 0 | 0 | 0 | 0 |
| **United Kingdom** | GBR | 0 | 2 | 0 | 0 | 1 | 0 | 0 | 0 | 0 | 0 | 0 | 0 | 0 | 0 |
| **United States of America** | USA | 0 | 0 | 0 | 45 | 457 | 59 | 0 | 0 | 0 | 0 | 0 | 0 | 7 | 12 |
| **Zambia** | ZMB | 11 | 0 | 0 | 0 | 0 | 0 | 0 | 0 | 0 | 0 | 22 | 0 | 0 | 0 |
| **Zimbabwe** | ZWE | 0 | 0 | 0 | 0 | 0 | 0 | 0 | 0 | 0 | 0 | 4 | 0 | 0 | 0 |

**S Table 2**. Feature coverage in the BRT models including a range of numeric variables as well as short name used for visualization purposes.

| **Variable** | **Short name** | **Coverage** | **Range** |
| --- | --- | --- | --- |
| TBV outcome | NA | 1.00 | NA |
| log(Livestock sum) | Livestock Sum | 1.00 | 0-19 |
| log(Human Pop Counts) | Pop Counts | 1.00 | 7-21 |
| log(Land Area sq km) | Land Area | 1.00 | 1-17 |
| IVSA Chapter Present | IVSA Chapter Present | 1.00 | NA |
| Total Ecoregions | NA | 1.00 | 1-62 |
| Koeppen-Geiger Climate Name | Climate Zone | 1.00 | NA |
| Pesticide Imports (USD$ million) | Pesticide Imports | 0.99 | 0-370 |
| Social Vulnerability | NA | 0.98 | 0.3-100 |
| Habitat Fragmentation | NA | 0.98 | 0.19-6 |
| % of Population with Access to Electricity | Electricity Access | 0.93 | 11-100 |
| Education Expenditures (% of GNI) | Education Spending | 0.92 | 0.4-24 |
| Livestock Production Index | Livestock Production | 0.90 | 36-190 |
| Annual People Treated for NTDs | NTD Treatment | 0.89 | 0-20 |
| Health Expenditures (% of GNI) | Health Spending | 0.88 | 2-16 |
| % of Population in Urban Areas | Urban Pop | 0.82 | 14-100 |
| % of Land Forested | Forested | 0.82 | 0-97 |
| % of Population Employed in Ag | Employed in Ag | 0.81 | 0.1-86 |
| Conflict Exposure (% of Pop Exposed) | Conflict (% of pop) | 0.81 | 0-100 |
| Conflict Exposure (tota events) | Conflict (total events) | 0.81 | 0-12 |
| Gini Index | NA | 0.78 | 25-62 |
| Adult Literacy Rate | Adult Literacy | 0.71 | 26-99 |
| % of Population Below National Poverty Line | Below Poverty Line | 0.60 | 1-82 |
| % of tertiary graduates from Ag, Vet, or Forestry progams | Vet/Ag/Forest Grads | 0.59 | 0-13 |

**S Table 3**. Additional covariate source information including spatial and temporal scale of variable. URL of the most comprehensive description of the variable is included. Gridded data (i.e., reported in km) was then averaged for within the country boundary from `rnaturalearth` package.

| **Driver type** | **Variable** | **Spatial** | **Temporal** | **url** |
| --- | --- | --- | --- | --- |
| Environmental | Total number of ecoregions | Polygons totaled up per country | Based on the year 1995 | https://databasin.org/datasets/e825e5a98fa14afb9a8bddeed0c47fc6/ |
| Environmental | Name of Köppen-Geiger Climate classification (mode) | The mode polygons within country boundary was selected | Represents the conditions between 1986-2010 | https://koeppen-geiger.vu-wien.ac.at/ |
| Environmental | Vegetation heterogeneity Index | 5 km | 2005 (single time period) | https://www.earthenv.org/texture |
| Environmental | % of land forested | country | 2023 or most recent reported year | https://data.worldbank.org/indicator/AG.LND.FRST.ZS |
| Exposure | Livestock Production Index | 1 km | 2015 (single time period) | https://data.worldbank.org/indicator/AG.PRD.LVSK.XD |
| Exposure | Livestock density | country | 2023 or most recent reported year | https://dataverse.harvard.edu/dataset.xhtml?persistentId=doi:10.7910/DVN/LHBICE |
| Exposure | % of population employed in agriculture | country | 2015 | https://data.worldbank.org/indicator/SL.AGR.EMPL.ZS |
| Knowledge | Pesticide imports | country | 2015-2017 (average) | https://data.worldbank.org/indicator/BM.AG.PEST.CD |
| Knowledge | Presence of an International Veterinary Students Association (IVSA) Chapter | 20 km | 2010-2020 (single time period) | https://www.ivsa.org/membership-directory/corporate |
| Knowledge | % of tertiary graduates from Ag, Vet, or Forestry schools | country | 2023 or most recent reported year | https://data.worldbank.org/indicator/SE.TER.GRAD.AG.ZS |
| Knowledge | Adult Literacy Rate | country | 2023 or most recent reported year | https://data.worldbank.org/indicator/SE.ADT.LITR.ZS?view=chart |
| Knowledge | Education expenditures | country | 2023, most recent reported year | https://data.worldbank.org/indicator/SE.XPD.TOTL.GB.ZS |
| Health Care | Health care expenditures | country | 1997-2023 (average) | https://data.worldbank.org/indicator/SH.XPD.CHEX.GD.ZS |
| Health Care | Reported average of annual people being treated for NTDs | country | 2015 |  |
| Wealth | Gini Index | 5 km | 2010-2020 (average) | https://data.worldbank.org/indicator/SI.POV.GINI |
| Wealth | % of population living below national poverty line | country | 2023, most recent reported year | https://data.worldbank.org/indicator/SI.POV.NAHC?view=chart |
| Wealth | Social Vulnerability Index | 1 km aggregated to 20 km |  | https://sedac.ciesin.columbia.edu/data/set/povmap-grdi-v1 |
| Wealth | % of population with access to electricity | country | 2023, most recent reported year | https://data.worldbank.org/indicator/EG.ELC.ACCS.ZS?view=chart |
| Reporting | Human population counts | 5 km | 2015 | https://sedac.ciesin.columbia.edu/data/set/gpw-v4-population-count-rev11/data-download |
| Reporting | Land area (sq km) | country | 2023, most recent reported year | https://data.worldbank.org/indicator/AG.LND.TOTL.K2 |
| Reporting | Conflict exposure (total events) | country | 2020-2023  (summed) | https://acleddata.com/conflict-exposure/ |
| Reporting | Conflict exposure (% of population exposed) | country | 2020-2023  (summed) | https://acleddata.com/conflict-exposure/ |
| Reporting | % of population in urban areas | country | 2023, most recent reported year | https://data.worldbank.org/indicator/SP.URB.TOTL.IN.ZS |

**S Table 4**. Variable importance rank (1-23) with relative influence (rel.inf), relative standard error (rse), and relative variance (rvar).

| **Rank** | **Variable** | **rel.inf** | **rse** | **rvar** |
| --- | --- | --- | --- | --- |
| **1** | Koeppen-Geiger Climate Name | 23.73 | 0.38 | 14.33 |
| **2** | Gini Index | 10.97 | 0.33 | 11.10 |
| **3** | log(Human Pop Counts) | 9.45 | 0.36 | 13.19 |
| **4** | Pesticide Imports (USD$) | 8.69 | 0.26 | 6.57 |
| **5** | log(Land Area sq km) | 7.92 | 0.30 | 8.97 |
| **6** | Livestock Production Index | 4.29 | 0.14 | 2.00 |
| **7** | Conflict Exposure (tota events) | 3.22 | 0.12 | 1.39 |
| **8** | % of Population Below National Poverty Line | 3.09 | 0.11 | 1.33 |
| **9** | IVSA Chapter Present | 2.84 | 0.21 | 4.39 |
| **10** | Conflict Exposure (% of Pop Exposed) | 2.69 | 0.11 | 1.26 |
| **11** | % of tertiary graduates from Ag, Vet, or Forestry programs | 2.50 | 0.10 | 0.97 |
| **12** | log(Livestock sum) | 2.35 | 0.10 | 1.00 |
| **13** | Adult Literacy Rate | 2.20 | 0.10 | 1.03 |
| **14** | % of Population in Urban Areas | 2.13 | 0.09 | 0.86 |
| **15** | Annual People Treated for NTDs | 2.13 | 0.08 | 0.72 |
| **16** | % of Population Employed in Ag | 2.09 | 0.09 | 0.79 |
| **17** | Health Expenditures (% of GNI) | 1.97 | 0.09 | 0.75 |
| **18** | % of Land Forested | 1.72 | 0.06 | 0.36 |
| **19** | Social Vulnerability | 1.61 | 0.09 | 0.85 |
| **20** | Total Ecoregions | 1.61 | 0.08 | 0.71 |
| **21** | Education Expenditures (% of GNI) | 1.43 | 0.06 | 0.31 |
| **22** | Habitat Fragmentation | 0.95 | 0.04 | 0.15 |
| **23** | % of Population with Access to Electricity | 0.42 | 0.02 | 0.06 |
